# CENTRE: A gradient boosting algorithm for Cell-type-specific ENhancer-Target pREdiction

**DOI:** 10.1101/2023.05.16.541035

**Authors:** Trisevgeni Rapakoulia, Sara Lopez Ruiz De Vargas, Persia Akbari Omgba, Verena Laupert, Igor Ulitsky, Martin Vingron

## Abstract

**Motivation:** Identifying target promoters of active enhancers is a crucial step for realizing gene regulation and deciphering phenotypes and diseases. Up to now, several computational methods were developed to predict enhancer gene interactions but they require either many epigenomic and transcriptomic experimental assays to generate cell-type-specific predictions or a single experiment applied to a large cohort of cell types to extract correlations between activities of regulatory elements. Thus, inferring cell-type-specific enhancer gene interactions in unstudied or poorly annotated cell types becomes a laborious and costly task.

**Results:** Here, we aim to infer cell-type-specific enhancer target interactions, using minimal experimental input. We introduce CENTRE, a machine learning framework that predicts enhancer target interactions in a cell-type-specific manner, using only gene expression and ChIP-seq data for three histone modifications for the cell type of interest. CENTRE exploits the wealth of available datasets and extracts cell-type agnostic statistics to complement the cell-type specific information. CENTRE is thoroughly tested across many datasets and cell types and achieves equivalent or superior performance than existing algorithms that require massive experimental data.

**Availability:** CENTRE’s open source code is available at GitHub via https://github.com/slrvv/CENTRE

## Introduction

Promoters and enhancers are the two major cis-regulatory elements that control the context-dependent gene transcription. Eukaryotic enhancers are bound by various transcription factors (TFs) and when activated they upregulate the expression of target genes by forming chromatin loops with their target promoters (Furlong and Levine, 2018). It is estimated that over a million potential enhancers exist in the human genome (ENCODE Project Consortium, 2012), vastly outnumbering human genes. Such significant redundancy shows that the activity of the enhancers is highly specific; only a subset of enhancers is active in a given cell type and orchestrates the lineage-specific gene expression (Visel *et al*., 2009).

As more mutations and genomic alterations of the non-coding genome become associated with regulatory elements, identifying gene targets of active enhancers is crucial for deciphering diseases and other phenotypes. Experimental methods such as Hi-C (Lieberman-Aiden *et al*., 2009), ChIA-PET (Fullwood *et al*., 2009), HiChIP (Mumbach *et al*., 2016), and Capture Hi-C (Mifsud *et al*., 2015) have revealed that chromatin architecture plays an important role in gene transcriptional regulation. When the resolution of these assays is high enough, these techniques can reveal individual Enhancer-Target (ET) contacts. However, high-resolution genome-wide loop data are only available for a limited number of human tissues/cell types and conditions. Further limitations such as low sensitivity, high cost, and technical challenges in loop-calling methods make capture-based techniques difficult to be widely applicable in EP identification (McCord *et al*., 2020), (Xu *et al*., 2020).

Several computational methods for identifying ET interactions have emerged. Correlating genomic and epigenomic signals at enhancers and promoters across multiple biosamples is the most common practice to detect ET pairs (Thurman *et al*., 2012; Sheffield *et al*., 2013). Although intuitive, these methods lack cell type (CT)-specific predictions while requiring a vast number of biosamples.

Supervised machine-learning methods train statistical models on sets of known interacting and non-interacting ET pairs annotated with various genomic, epigenomic, and transcriptomic features. Once the model is trained in one or several cell types, in principle, it could predict ET pairs in any other cell type. One constraint of these methods is that they require multiple experimental assays to generate CT-specific ET predictions. TargetFinder (Whalen *et al*., 2016) integrates hundreds of cell-specific genomics datasets to annotate ET pairs, making the method applicable only to a few rich annotated cell lines. Inflated performance due to dependencies in training and test datasets is another limitation in assessment of the supervised learning models (Cao and Fullwood, 2019). An unbiased and robust computational approach that can be easily applied to any cell type while it claims for a feasible number of experiments is still missing.

Given a hitherto less studied tissue or cell type, the challenge lies in predicting its ET interactions using a minimum of experimentally derived information. We here present Cell-specific ENhancer Target pREdiction (CENTRE) which requires for the cell type under study only RNA-seq results and ChIP-seq data for H3K27ac, H3K4me1, and H3K4me3. This cell type specific information is combined with correlations between enhancer and promoter regions across many cell types by means of a machine learning method, extreme gradient boosting. Through this combination of generic, across-cell-type information with cell type-specific information we obtain a prediction accuracy which is en par or better than established tools. At the same time the requirement for genomic data about the new cell type is low enough to allow for easy and routine use of CENTRE for predicting ET interactions in a new cell type.

## Results

### CENTRE algorithm

We designed a machine learning pipeline called CENTRE (Cell-specific ENhancer Target pREdiction), that predicts ET interactions in a CT-specific manner (Fig 1A). CENTRE builds on the ENCODE Registry of candidate cis-regulatory elements with enhancer-like signatures (cCRE-ELS) (ENCODE Project Consortium *et al*., 2020) and GENCODE transcription start sites (TSSs) (Wright *et al*., 2016). During training, the input consists of pairs of enhancers and promoters, labeled as “interacting” or “not interacting” as given in the BENGI ground truth dataset (Moore *et al*., 2020). The algorithm annotates potential ET interactions with a minimum set of CT-specific information coming from histone marks (HMs), gene expression, and generic information derived from ensembles of biosamples and distance. Then a pre-trained extreme gradient boosting classifier (Chen and Guestrin, 2016) computes a probability for an annotated ET pair to interact in the cell type of interest. We however limit the search for ET pairs to a distance of 500 KB which is a commonly accepted threshold for ET interactions (van Arensbergen *et al*., 2014) that matches the median size of topologically associated domains (TADs) (Dekker *et al*., 2013).

**Figure 1:**
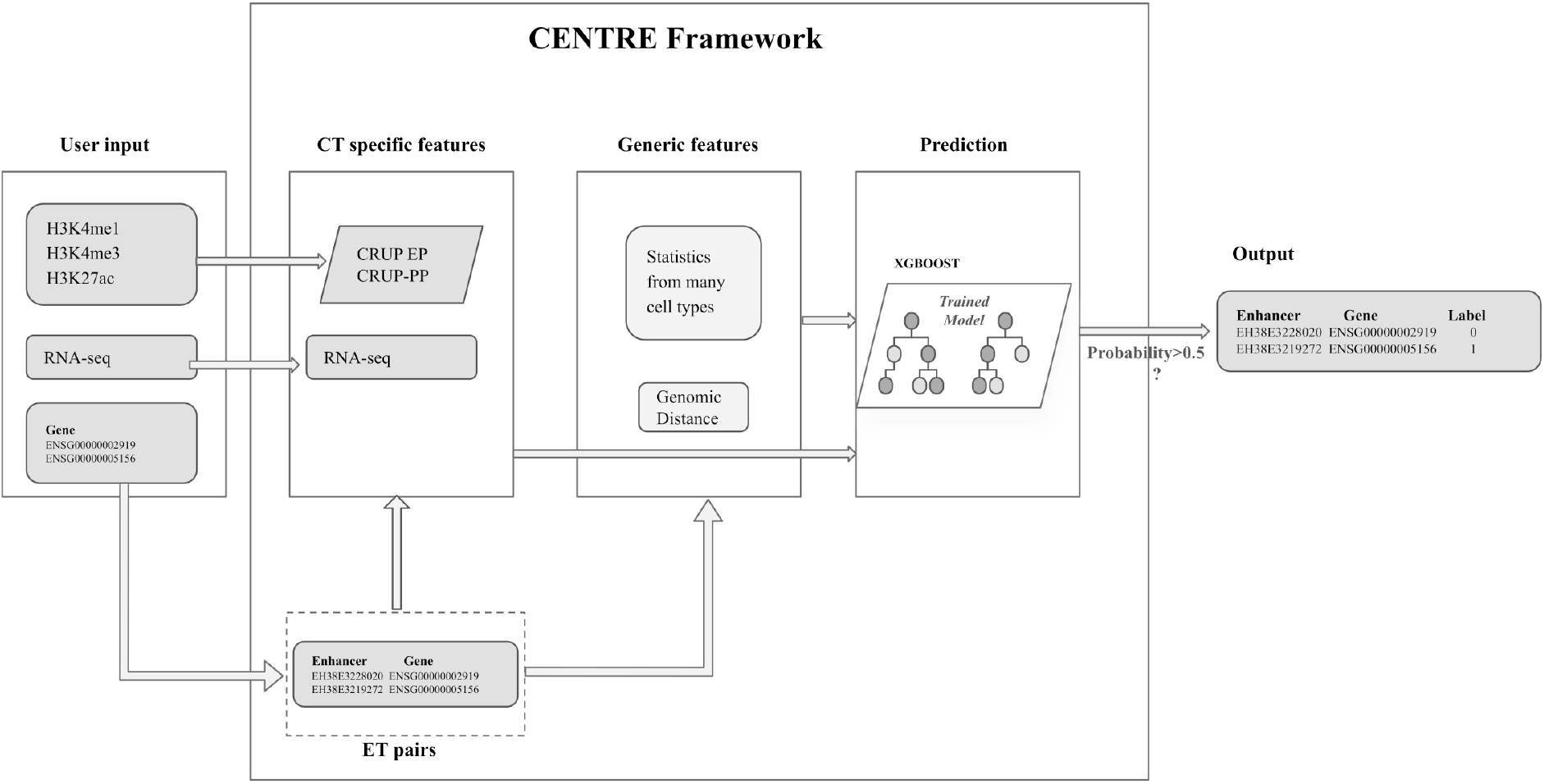
Outline of CENTRE framework: The user provides target genes of interest with CT specific RNA-seq and H3K27ac, H3K4me1 and H3K4me3 ChIP-seq data. CENTRE extracts all the cCRE-ELS within 500KB of target genes and computes CT-specific and generic features for all potential ET pairs. ET feature vectors are then fed to a pre-trained XGBOOST classifier, and a probability of an interaction is assigned to ET pairs. ET pairs with higher probability than 0.5 are labeled as interacting pairs.

For predicting ET pairs in a new cell type, our algorithm only requires RNA-seq data as well as ChIP-seq data of the histone modifications H3K27ac, H3K4me1 and H3K4me3 determined for this particular cell type. These histone modifications reflect the activity status of an enhancer or promoter, while the RNA-seq data inform the machine learning procedure about the transcriptional outcome of a possible ET interaction. The classifier is trained using this information for many available cell types in conjunction with generic, across-cell-type statistics of epigenetic and transcriptomic signals in enhancers and targets. We think of the statistics between regulatory elements across cell types as providing the potential for an interaction, while the CT specific information serves to predict whether an interaction is realized and leads to gene activation or upregulation in a particular cell type.

### Features reflecting generic, across-cell-type information

The generic, across-cell-type information is based on the rationale that when a gene’s expression is increased by an enhancer, then one expects to find this enhancer accessible, or more generally active, in those cell types where the gene is upregulated. Conversely, one expects the gene to be more lowly expressed when the enhancer is inactive. This logic suggests looking for correlations between enhancer accessibility and gene expression across many cell types, realized by many existing methods (Sheffield *et al*., 2013).

However, simply using correlation, e.g., of DNAse accessibility patterns with gene expression, does not suffice to point out significant ET interactions because of the CT-specific activity of enhancers. As an example, the TTC39C gene interacts with the distal enhancer-like element EH38E1904551 in the GM12878 cell line according to the ChIA-PET experiment targeting RNAPII (Tang *et al*., 2015). The expression of TTC39C across 112 tissues (Sheffield *et al*., 2013) though does not correlate with the accessibility of the EH38E1904551 element in the same tissues as can be seen in Fig. 2A. In this figure, for only a few tissues there is a clear enhancer accessibility signal as well as high expression, while generally the epigenetic signal is low despite the gene being highly expressed. These kinds of examples demonstrate why a correlation coefficient has a low predictive value identifying ET pairs.

**Figure 2:**
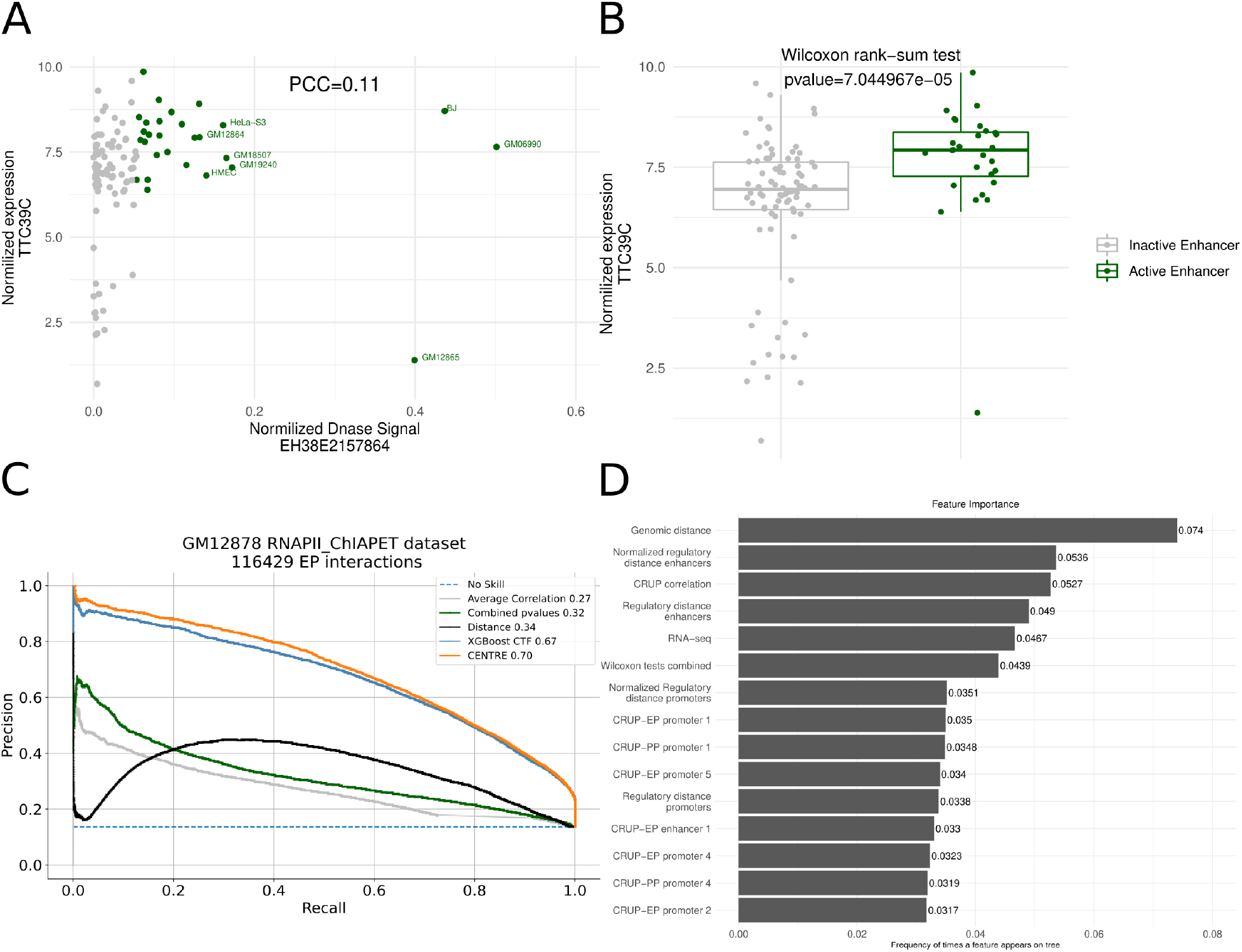
A) Scatter plot of normalized TTC39C expression and DNase signal at EH38E1904551 across 112 human cell types. Green dots represent cell types with higher accessibility (upper quantile DNase signal). Although TTC39C is expressed across many cell types, EH38E1904551 presents high DNase signals predominantly in lymphoblastoid cell lines, resulting in a Pearson Correlation Coefficient of only 0.11. B) Boxplots of TTC39C expression in cells where the EH38E1904551 is accessible (upper quantile DNase signal across 112 cells) versus cells where EH38E1904551 is not accessible. The significant p-value of the Wilcoxon rank sum test finer illustrates the strength of the ET interaction. C) Precision Recall Curve plots show the performance of individual feature sets on classifying the GM12878 RNAPII-ChIAPET dataset. The machine learning algorithm achieves its best performance when trained on CT specific features and generic evidence of the ET interactions. The performance of the gradient boosting algorithms is measured after 12-fold cross validation, training and testing in different chromosomes. CENTRE was optimized on the GM12878 RNAPII-ChIAPET data using a nested cross validation scheme (Methods). D) Ranking of feature importance, based on the relative number of times a particular feature occurs in XGBoost trees (feature weight over weights of all features).

To mitigate this situation, we draw on another statistic. We divide the cell types according to present or absent enhancer activity. This allows comparing gene activity when the enhancer is active vs. inactive (Methods) by performing a Wilcoxon rank-sum test between target transcriptional signals between cells where the enhancer is active and those where it is not active. We use its associated p-value as an indicator of the significance of the ET interaction. For the case of accessibility as a descriptor of enhancer activity, this is visualized in Fig. 2B where one can see that in those cell types where the enhancer under study is active the target gene tends to be more highly expressed, resulting in a significant p-value of the Wilcoxon rank-sum test.

We further extend this approach by including other descriptors of enhancer activity for cCREs and GENCODE TSS activity. Descriptors for enhancer activity comprise DNAse accessibility and eRNA expression as measured by CAGE tags. On the side of the target promoter/gene activity we use DNAse accessibility and downstream transcript level as measured by microarrays and CAGE-seq. We use three publicly available datasets (Thurman *et al*., 2012), (Sheffield *et al*., 2013), and (Andersson *et al*., 2014) measuring the enhancer and target aforementioned signals in an ensemble of cell types and we calculate the Wilcoxon rank-sum test p-value for all enhancers and target pairs within 500KB. In addition we employ enhancer probabilities generated by CRUP-EP (Ramisch *et al*., 2019) which uses H3K4me1, H3K4me3, and H3K27ac ChIP-seq data and predicts the enhancer activity of a genomic region. We used a dataset consisting of CRUP-EP probabilities and RNA-seq TPM values across 66 ENCODE cell types (Supplementary Table 1), and applied the Wilcoxon rank-sum test analysis. For annotating ET pairs, we summarize these four p-values into a single Fisher’s combined probability (Methods).

Based on the way CRUP works, we could also extract probabilities that a region constitutes an active promoter, namely CRUP-PP promoter probability (Methods). We downloaded H3K4me1, H3K4me3 and H3K27ac ChIP-seq data for 104 cell types from the ENCODE portal (Supplementary Tables 2, 3, & 4), and we applied the CRUP-EP and CRUP-PP functions. We complement the across-cell-type information with the Pearson’s correlation coefficient between CRUP-EP and CRUP-PP probabilities over the 104 cell types (Methods). Thus, the generic information of CENTRE results in three feature values that will be input to the machine learning classifier: 1) Fisher’s combined probability, 2) correlation coefficient of CRUP probabilities and 3) the genomic distance between the ET pairs.

### Features reflecting cell type-specific information

Aggregate statistics obtained from many biosamples are insufficient to delineate CT-specific links. Thus, computational ET prediction methods also use CT-specific assays to capture cell type dependent interactions. We designed two kinds of features. Firstly, we want to capture whether an enhancer and promoter are each active. To this end, we again apply the CRUP-EP and CRUP-PP functions (Methods) to compute from histone marks (HMs) the probabilities for cCRE-ELS and TSS to be active as enhancers or promoters, respectively. Additionally, we use the expression level of the respective target gene as given by the CT-specific RNA-seq data.

A very important feature in predicting whether cCRE-ELS might target a certain promoter is the genomic distance between the regulatory elements. However, this is a generic feature carrying no information about a particular cell type. We again exploit CRUP predictions for regulatory elements to upgrade the genomic distance feature to a “regulatory distance” which describes whether there are many active regulatory elements in the genomic region between the ET pairs, or whether that region is largely devoid of regulatory activity. We apply CRUP-EP and CRUP-PP to the window between the two regulatory elements and extract the fraction of regions classified as enhancers or promoters, respectively. Note that regulatory distance (RD), in contrast to genomic distance alone, captures CT-specific information.

### Integrating features into a machine learning framework

Taken together we form features from the following information: 1) Statistics among regulatory element activities in many cell types, as described by the Wilcoxon rank sum test and Pearson’s correlation coefficient, 2) genomic distance and regulatory distance and 3) CT-specific regulatory element activity predictions obtained using CRUP and gene expression. The CENTRE algorithm combines all these orthogonal sources of information into a single probability score describing the likelihood that an enhancer targets a particular TSS in a given cell type. While we exploit ample information to create a priori features from association signals across many cell types, the CT-specific information needed comprises only gene expression plus the three HMs, namely H3K4me1, H3K4me3 and H3K27ac. This latter information is widely available for many cells and can be relatively easily collected for a cell type which was not profiled yet. A single feature vector is generated from the combination of across-CT and within-CT-specific information which feeds an XGboost classifier (Chen and Guestrin, 2016).

For a proof of concept we tested the ability of individual features to correctly classify ET interactions as derived from GM12878 RNAPII-ChIAPET data (Tang *et al*., 2015) and labeled by (Moore *et al*., 2020). In Fig. 2C, we can clearly notice the advantage of the Fisher’s combined probability extracted from three publicly available datasets over the average Pearson correlation derived from the same datasets. However, both generic features perform poorly on distinguishing true and false ET interactions on the GM12878 cell line, being inferior to the baseline distance method. When we train the XGboost classifier with CT-specific CRUP probabilities on regulatory elements as well as on the window between them (RD), together with the GM12878 RNA-seq data the performance significantly increases. Combining all the orthogonal variates (across cell-type information, genomic distance, CT-specific gene expression, CRUP predictions) into a single feature vector, CENTRE achieves the highest area under precision recall curve (AUPRC), while still using few CT-specific features.

### Feature importance

We ranked features according to their importance, using the number of times they appear in XGBoost trees. Among the leading features in Fig. 2D one finds both generic and CT specific features. Genomic distance (position 1), CRUP correlation (position 3), and Wilcoxon tests combined (position 6) are generic features, while positions 2, 4 and 5 are occupied by CT specific features. This underlines that both generic and CT-specific features contribute to the overall performance.

With versions of RD scoring high in feature importance, we further checked whether regulatory distance is in fact more informative than mere genomic distance. To this end, we collected enhancer-promoter pairs which interact in LCLs but not in HeLa cells, based on the annotation provided by (Moore *et al*., 2020). Clearly for all these ET pairs the genomic distance remains constant independent of cell type. Fig. 3A shows a scatter plot comparing enhancer RD (Normalized CRUP-EP in the window) between LCLs and HeLa cells. The regression line has a smaller slope than the identity indicating that enhancer RD is smaller in the LCLs where the pairs actually interact. Another version of RD estimates the number of active promoters between two regions. This feature, termed Normalized CRUP-PP in the window, is plotted in Fig. 3B which shows that this count is dramatically lower in the LCLs as compared to HeLa cells.

**Figure 3:**
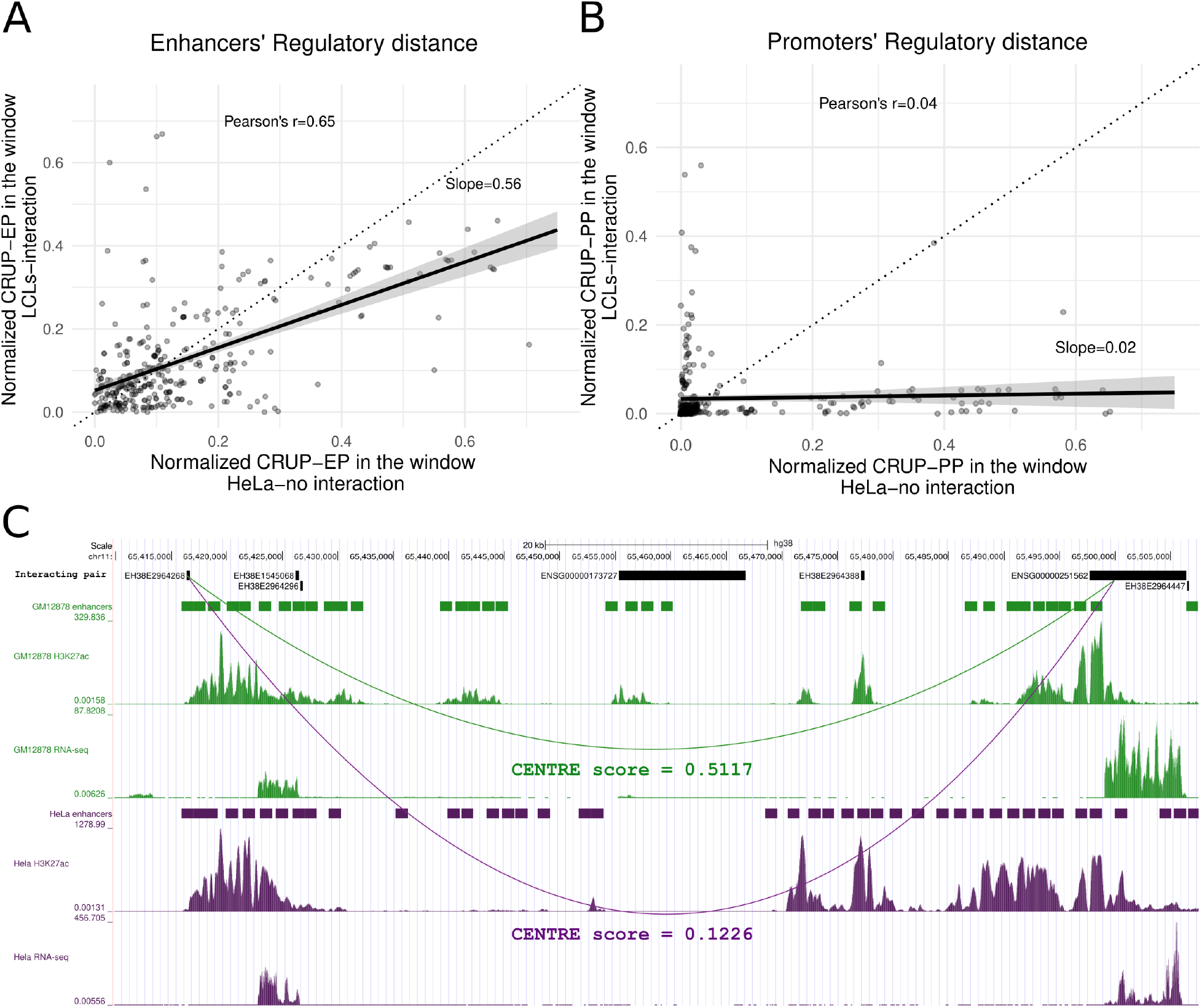
A) Scatterplot of enhancers’ RD, captured by the normalized CRUP-EP scores applied to the window between 334 ET pairs that interact in LCLs but not in HeLa cell line. B) Scatterplot of promoters’ RD, captured by the normalized CRUP-PP scores applied to the window between 334 ET pairs that interact in LCLs but not in HeLa cell line. We can notice that both enhancers and promoters’ RD tend to be smaller in LCLs than HeLa cells. C) Genome Browser view of the MALAT1 gene (ENSG00000251562) which is targeted by the upstream CRE EH38E2964268 in GM12878 but not in HeLa. The enhancers’ RD is smaller in GM12878 compared to HeLa, which is also confirmed by the H3K27ac signals. CENTRE correctly uncovers the CT specific true interaction in GM12878, while it assigns a very small probability in HeLa, where the specific ET pair does not interact.

An example of an ET pair that does not have the same RD in cells where there is true regulation is given in Figure 3C. The cCRE-ELS EH38E2964268 targets the metastasis associated lung adenocarcinoma non-coding RNA gene MALAT1 (ENSG00000251562) in the GM12878 cell line but not in HeLa. As we can notice the enhancer’s RD between the specific ET pair is smaller in GM12878 than HeLa, as indicated by CT-specific CRUP-EPs and the H3K27ac track, as well as RNA-seq signals. CENTRE algorithm correctly predicts the CT specific link, assigning a probability of interaction of 0.51 in GM12878 but a lower probability of 0.12 in the same pair in HeLa.

### Validation using BENGI dataset

For the validation of CENTRE on multiple cell types, we used the *Benchmark of candidate Enhancer-Gene Interactions* (BENGI) established by (Moore *et al*., 2020). In their paper the authors evaluated several computational enhancer target identification methods including, in particular, TargetFinder (Whalen *et al*., 2016) which was shown to be the best-performing method across all datasets (Moore *et al*., 2020). For the purpose of evaluation, the study puts together a comprehensive testbed annotating pairs of cCREs-ELS and GENCODE TSS with experimental evidence derived from either 3D chromatin interactions (ChIa-PET), HiC, genetic interactions, or CRISPR/dCAS9 perturbations for 13 cell types. To avoid evaluation bias due to dependencies between training and test datasets, the authors also provide 12 cross-validation (CV) groups split by chromosome. This ensures that testing is always performed in different genomic regions than training. We use BENGI datasets and follow their suggested routine of 12-fold CV such that our results should be fully comparable to results reported by Moore et al. 2020. For evaluation we use F1-score. F1-score is a well-suited metric for highly imbalanced datasets as is the case with the few reported positive interactions in comparison to a large number of non-interacting pairs.

The full TargetFinder model uses roughly 101 epigenomic and transcriptomic experiments from histone modification ChIP-seq, TF ChIP-seq, DNase-seq, and CAGE-seq, yielding 303 features. Our CENTRE method computes 28 features stemming from only four experiments, namely three ChIP-Seq histone marks and RNA-seq. For the initial comparison we used ET datasets derived from five commonly used cell types where all experiments required by TargetFinder are available, which then also includes the four experiments used by CENTRE. Figure 4A shows the performance of CENTRE compared to TargetFinder (at 12-fold CV) in terms of F1-score. CENTRE achieves a higher F1-score than TargetFinder in 11 out of 13 benchmark datasets, where positive pairs were extracted from five experimental assays applied in five cell lines. Especially when positive pairs come from Hi-C loops, both methods’ performance is limited. However, CENTRE is more efficient in uncovering ET interactions across many cell types and experimental techniques. TargetFinder in contrast, although requiring substantially more CT specific information, performs better than CENTRE in only two ChIAPET RNAPII datasets.

**Figure 4:**
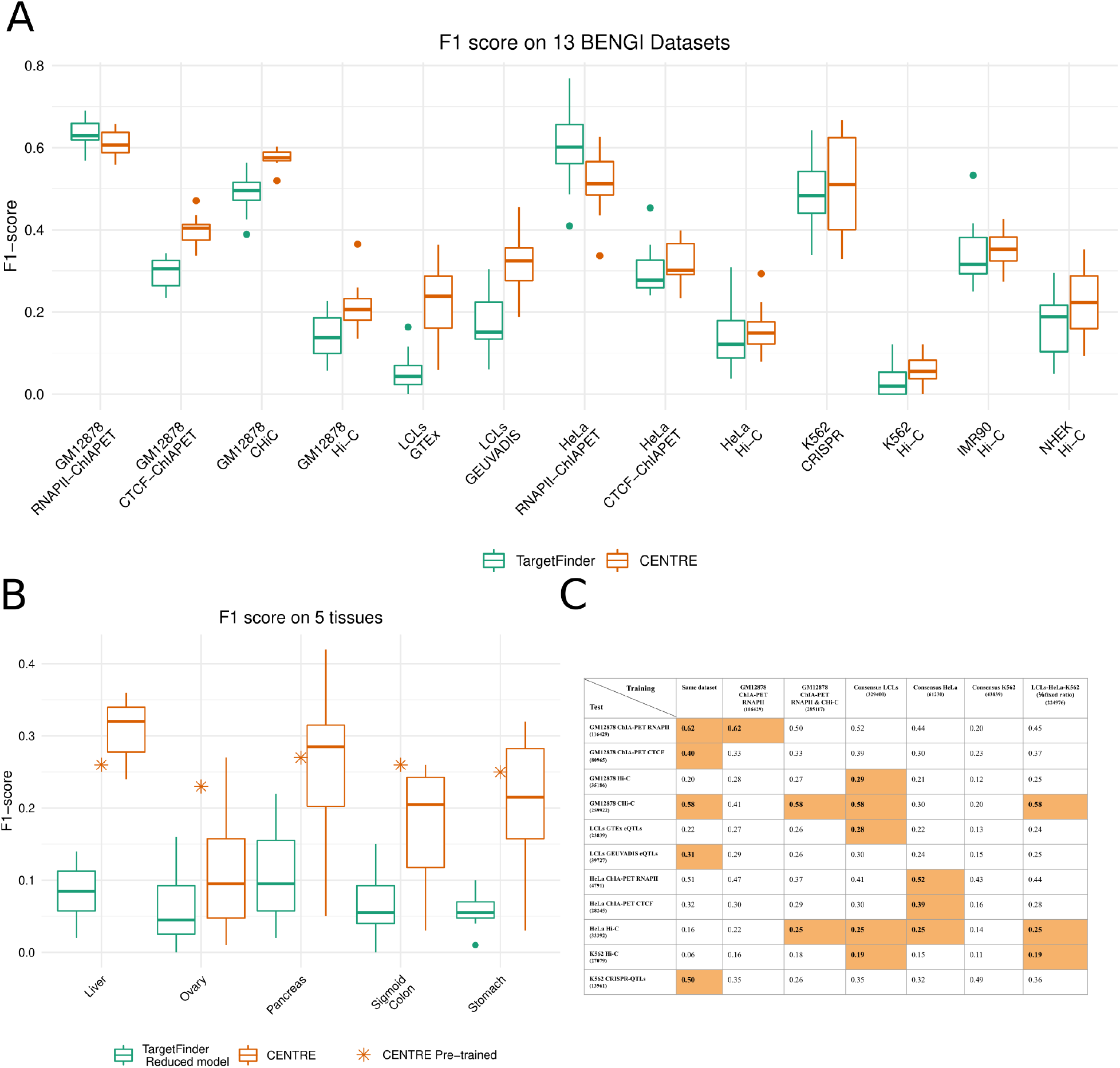
A) Boxplots representing the F1 scores after 12-fold CV achieved by CENTRE and TargetFinder applied in 13 BENGI datasets (five cell lines) where all experimental assays required by TargetFinder are available. B) Comparison of CENTRE and TargetFinder in five tissue datasets where a subset of experimental assays required by TargetFinder was used (TargetFinder Reduced model). The asterisk denotes the F1 score of the pre-trained CENTRE on the consensus LCL dataset and applied on the five tissue datasets. C) Mean F1 scores after 12-fold CV of CENTRE when trained and tested on different datasets. For each test dataset, orange cells highlight the best performing training dataset.

We extended our evaluation in five tissue datasets where the positive pairs were extracted from eQTL mapping. Since not all TF ChIP-seq datasets needed for training of TargetFinder were available, we reimplemented a reduced TargetFinder model using a subset of 13 features coming from DNase, H3K4me3, H3K27ac, and CTCF experimental assays and genomic distance, as suggested by (Moore *et al*., 2020). Figure 4B shows that CENTRE’s advantage becomes even more evident in these five tissues, significantly outperforming the reduced TargetFinder model. Even though the reduced TargetFinder model uses more CT specific information than CENTRE, still CENTRE predicts CT specific ET pairs correctly more often, relying only on a minimal amount of CT specific information.

### Pre-trained CENTRE Framework

Training a classifier is a complicated process and rather than requiring the user to train a classifier a pre-trained classifier should be provided. Therefore, we provide a pre-trained classifier software that we have trained on a data set of known ET interactions and that is then ready to make new ET predictions in any cell type and context. To this end, the selection of the training dataset is a critical factor for the algorithm’s ability in future predictions. Based on the information summarized in Figure 4, we focused on K562, GM12878, HeLa cell lines. For these cell types, training ET links were derived from multiple experimental assays in the BENGI benchmark collection. Based on the expectation that these experimental methods capture different aspects of ET interactions, we created joint cell-line-specific consensus datasets and a comprehensive composite dataset consisting of all labeled ET interactions from the three cell lines in a fixed positive-to-negative ratio. We also combined the two best performing datasets according to Figures 4A & B (GM12878 RNAPII-ChIAPET and CHi-C, Figure 4A) and kept intact the GM12878 RNAPII-ChIAPET (Tang *et al*., 2015) dataset in the training collection since CENTRE achieved its best performance when applied to it.

For the classifiers trained on this information, we performed extensive performance evaluation on 11 ET datasets coming from three cell lines and six different experimental assays, using the 12 CV scheme to avoid overfitting. According to the results shown in figure 4C, when CENTRE is trained on the consensus LCL training dataset, it presents the most solid performance, achieving the best F1-score in five out of 11 testing datasets and the second-best in another three datasets. Noteworthy, among its best-performing test sets, are the K562 and HeLa datasets coming from Hi-C loops, displaying good performance across cell-types.

Once we homed in on the training set, CENTRE was trained and optimized on the whole consensus LCL dataset (Methods). As a proof of concept, we applied the pre-trained classifier on the five eQTL tissue datasets of Fig. 4B. As we can notice the pre-trained algorithm on the consensus LCL dataset and applied to the whole five tissue datasets performs similarly to the method when trained and tested on the same datasets with 12-fold CV. Noteworthy, it achieves an even better f1 score than the median 12-fold CV scores for three out of five tissue datasets, showing its predictive ability when applied to different cell types. We provide the pre-trained CENTRE framework as ready-to-use R software. For application to a particular cell type, it takes as input the CT specific H3K4me1, H3K4me3, H3K27ac, ChIP-seq data, and RNA-seq TPM values. Then, for a user provided target gene of interest the software predicts the interacting cCRE-ELSs. There is no need for retraining and the inclusion of all the other sources of information in the training is invisible to the user.

## Discussion

In this work we have put forward the CENTRE method to predict interacting enhancer-promoter pairs in a cell type of interest. In a real-life application, that cell type of interest will typically be a less studied cell type and our method requires only a limited set of experimental data to base the prediction on. When training the method, however, we include a wealth of available data to establish a space of feasible interactions. These interactions get reweighted according to the cell type specific information. This process is transparent to the user who does not need to retrain the program but instead only provides the cell type specific data to the trained algorithm. Despite this simplicity in applying CENTRE, the quality of its predictions is generally comparable to, and in some cases better, than the best existing methods.

Designing a machine learning method not only yields the benefit of the final product, the program, but also allows to study which features contribute and improve the quality of the predictions. We did not attempt to assemble large numbers of features but put a lot of effort into the careful design of the features. In this process we experienced a few surprises. Firstly, in the context of distilling the information from available data across cell types, we found that a rank-sum test among two regulatory features of many cell types adds valuable information on top of that provided by correlations. One is tempted to speculate that enhancer and promoter activity across many cell types need not be highly correlated but rather they only display similar ranking, i.e., when the enhancer-promoter link is active one picks up a signal and when the link is inactive there is no correlation of activities.

The other insight from the feature design concerns the regulatory distance. While genomic distance clearly plays an important role in enhancer-promoter interaction, regulatory distance describes how much regulatory activity there is in the interval between enhancer and promoter in the cell type under study. We have shown that inclusion of this feature improves prediction, and that regulatory distance is connected to the probability of an ET link. This suggests that for the cell it is easier to establish a specific chromatin interaction when in the loop that is excluded there is little other regulatory activity.

Although CENTRE performs well and compares favorably with TargetFinder, the F1 score on many datasets is still low. To a certain degree this simply reflects the inherent difficulty of the problem of enhancer target prediction. We also tried to put our method to the toughest tests and present fair comparisons and results, avoiding overfitting issues due to dependent genomic regions in training and test sets. Still, even the choice of validation data strongly influences any performance measurement. We notice that the Hi-C datasets consistently exhibited the lowest overall performance. One possible reason is that Hi-C maps even at 5kb resolution are too coarse and cannot be used to link distal regulatory elements to their target genes (Zhang *et al*., 2018). Another limitation stems from technical challenges in calling Hi-C loops, where different loop-calling methods can produce markedly different results (Forcato *et al*., 2017).

Given the difficulty of the enhancer target prediction problem, there is clearly room for future improvement of our method. In the current version we derive the space of feasible interactions from an analysis of large amounts of epigenetic data. Here the question is whether and how to capitalize on Hi-C data when it is available. Also, we still have to limit the predicted interactions to a distance of 500 kb to avoid large numbers of false positives. Future methodological improvements will hopefully allow extending this interval.

The pre-trained CENTRE framework is provided as ready-to-use R software where the user gives target genes along with cell type-specific H3K4me1, H3K4me3, H3K27ac, ChIP-seq, and RNA-seq TPM data and receives all the interacting and non-interacting predicted enhancers for the genes of interest. Thus, CENTRE provides an accurate, pragmatic framework to distinguish genomic interactions without the need for extensive and costly experiments.

## Methods

### Processing of BENGI datasets

We used the All-Pairs.Natural-Ratio BENGI collection of datasets for the training and evaluation of CENTRE. BENGI uses the hg19 annotation of cCREs-ELS and GENCODE v19 TSS. We updated all ET pairs of the BENGI datasets on the hg38 annotation using the UCSC Genome Browser liftover facility (Kent, 2002). Then we overlapped the uplifted regions with the latest version 2 of ENCODE Registry of cCREs on hg38 (findOverlaps function from GenomicRanges library (Lawrence *et al*., 2013)). Regarding the targets, we used the basic gene annotation of GENCODE Release 40 (GRCh38). The hg38 Registry of cCREs-ELS has a variable size between 150-350 bp (median length 286 bp), while the hg19 annotation has a variable size between 50-16,633 bp (median length 352 bp). In case of a hg19 cCREs-ELS overlapping more than one hg38 cCREs-ELS, multiple ET pairs were created with the mapped hg38 cCREs-ELS and the GENCODE TSS assigning the original label. We kept only ET pairs that are located within 500KB. We downloaded RNA-seq TPM values for the 13 biosamples from ENCODE (Supplementary Table 1). The positive interactions were further processed so that the interacting gene had a TPM value greater than zero. We kept negative interactions with either a matching TSS or a matching cCREs-ELS in the positive set. All the processed BENGI datasets used in this study can be found at: http://owww.molgen.mpg.de/~CENTRE_data/BENGI_processed_datasets.zip

### CT specific features

#### CRUP-EP probabilities on cCREs

ENCODE cCREs-ELS, and GENCODE TSSs were extended on both sides to have a length of 500 bp. We downloaded H3K4me1, H3K4me3, and H3K27ac ChIP-seq data from the ENCODE portal (Supplementary Tables 2,3,& 4) for all the cell types examined in the study, and we applied the CRUP-EP algorithm. CRUP-EP outputs the enhancer probability for every 100-bp genomic bin. We overlapped the obtained probabilities with the 500 bp enhancer and target regions (findOverlaps function from GenomicRanges library (Lawrence *et al*., 2013), resulting in five enhancer probabilities scores for the enhancer regions and five for the target regions.

#### CRUP-PP probabilities on cCREs

CRUP-EP uses a combination of two binary random forest classifiers and assigns enhancer probabilities to each 100-bp bin (Eq. 1). The first classifier discriminates between active genomic regions (active promoters, enhancers) and inactive genomic regions (inactive promoters, remaining intra- and intergenic regions). The second classifier distinguishes enhancers from promoters, given that the bin is active.

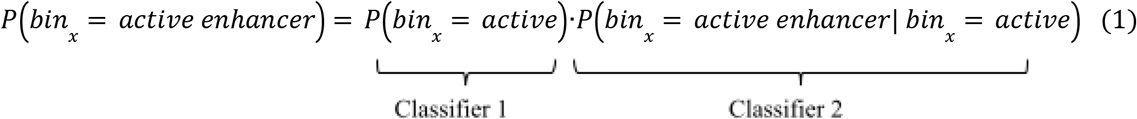

We used the complementary probability of the second classifier that distinguishes enhancers from promoters such that it can return the probability of the bin to be an active promoter, given that the bin is active. (Eq. 2). We call the output CRUP Promoter Probability (CRUP-PP).

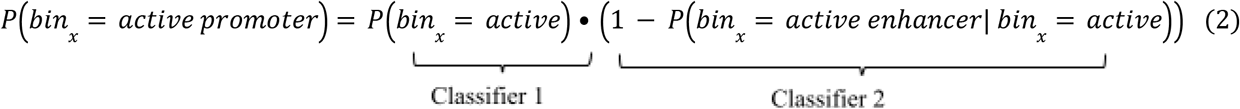

We overlapped the obtained probabilities with the 500 bp enhancer and target regions (findOverlaps function from GenomicRanges library (Lawrence *et al*., 2013)), resulting in five promoter probabilities scores for the enhancer regions and five for the target regions.

#### Regulatory Distance

We extracted the CRUP-EP and CRUP-PP activity scores for the window between the ET pair. RD features consist of four different values:1) the number of bins where CRUP-EP>0.5, 2) the number of bins where CRUP-EP>0.5 divided by the total number of bins in the ET window, 3) the number of bins where CRUP-PP>0.5, 4) the number of bins where CRUP-PP>0.5 divided by the total number of bins in the ET window.

#### RNA-seq

RNA-seq transcripts per million (TPM) values for all biosamples considered in the study were downloaded from the ENCODE portal (Supplementary Table 1).

### Generic Features

#### CAGE-seq dataset

We downloaded the RLE normalized expression TPM tables for enhancers and genes from the FANTOM5 portal (https://fantom.gsc.riken.jp/5/datafiles/reprocessed/hg38_latest/extra/). We averaged TPM values for the enhancers and for the genes across replicates resulting in 848 different cell types. We intersected cCRE-ELS with enhancer CAGE peaks (findOverlaps function from GenomicRanges library (Lawrence *et al*., 2013)). If a cCRE-ELS region overlapped with more than one CAGE-defined enhancer, we added the TPM values of the overlapped CAGE enhancers. We used the Wilcoxon rank-sum test to compare the gene expression in samples where the enhancer is active (TPM>0) and inactive (TPM=0).

#### DNAse-hypersensitive region dataset

We downloaded normalized counts across 112 cell types for DNase-hypersensitive sites or DHSs (dhs112_v3.bed) from http://big.databio.org/papers/RED/supplement/. We used UCSC liftOver (Kent, 2002) to obtain the hg38 coordinates of DHSs. We intersected cCRE-ELS and target regions with the DHSs (findOverlaps function from GenomicRanges library (Lawrence *et al*., 2013)). If cCRE-ELS and target regions overlapped with more than one DHS, we added the corresponding DHSs counts. We ranked cell types based on the enhancer DHSs normalized counts and selected the top quantile (0.25) as the ones with the higher enhancer activity. We then used the Wilcoxon rank-sum test to compare the target DHSs signal in samples where the enhancer has higher activity than the rest of the cell types.

#### DNAse-seq - gene expression dataset

We downloaded the normalized microarray gene expression for 112 cell types (exp112.bed) that match the DNAse-hypersensitive dataset from http://big.databio.org/papers/RED/supplement/. We used the processed DNA-seq dataset for the enhancer regions and we applied the Wilcoxon rank-sum test to compare the target gene expression in samples where the enhancer has higher DNAse activity (top quantile) than the rest of the cell types.

#### CRUP-EP-gene expression dataset

We downloaded H3K4me1, H3K4me3, H3K27ac, ChIP-seq data, and RNA-seq TPM values for 66 matched cell types from the ENCODE portal (Supplementary Tables 1-4). We applied the CRUP-EP function and extracted the enhancer probabilities for cCRE-ELS regions, averaging them across the five bins. If the average CRUP-EP probability was greater than 0.5, we considered the enhancer region as an active enhancer. We used the Wilcoxon rank-sum test to compare the gene expression TPM values in cell types where the enhancer predicted active and inactive.

#### Fisher’s combined probability

We combined the four Wilcoxon rank-sum test p-values into a single p-value using Fisher’s method (Statistical methods for research workers, 1935). We used the negative logarithm of the combined p-value as the final feature in our classification.

#### CRUP-EP & CRUP-PP correlation

We downloaded H3K4me1, H3K4me3, H3K27ac, ChIP-seq data for 104 cell types from the ENCODE portal (Supplementary Tables 2-4). We applied CRUP-EP and CRUP-PP functions in all cell types and extracted the CRUP-EP and CRUP-PP predictions for cCRE-ELS and target regions, respectively. We summed the probabilities over the five bin regions and computed the Pearson correlation coefficient across the 104 cell types.

#### Genomic distance

We computed the distance between the middle-point of cCREs-ELS and target TSSs according to the ENCODE Registry of cCREs on hg38 and the basic gene annotation of GENCODE Release 40, respectively. We used the absolute value of distance for the classification. All the features are given analytically in the Supplementary Table 6.

#### CENTRE algorithm

We applied the XGBoost (Chen and Guestrin, 2016) algorithm (python xgboost.XGBClassifier) with the logistic regression learning objective for binary classification. To control the unbalance of positive and negative samples we set scale_pos_weight=5. We used random_state=0 for reproducibility.

We initially optimized the algorithm on the GM12878 RNAPII-ChIAPET data (Tang *et al*., 2015) using GridSearchCV with a nested CV scheme for the model selection (inner folds=3, outer folds =12). Based on the average precision reported in the inner CV we selected the following parameters: colsample_bytree=0.7, gamma=1.0, learning_rate= 0.1, max_depth=5, n_estimators=300, reg_lambda=0, subsample=0.9. The outer CV is adopted from (Moore *et al*., 2020) to ensure that testing is always performed in different genomic regions than training—the results in Fig. 2C display only the outer CV performance where no hyperparameter tuning was carried out. We used the same parameters without further optimization in the rest of the BENGI datasets (Fig. 4A and 4B).

We finally optimized the XGBoost algorithm on the consensus LCL datasets, using the RandomizedSearchCV function to find the optimal parameters. We used the customized CV scheme suggested in (Moore *et al*., 2020) to avoid overfitting. Based on the f1 score performances we selected the following parameter set for the pre-trained CENTRE: colsample_bytree=0.7, gamma=0.25, learning_rate= 0.1, max_depth=10, n_estimators=300, reg_lambda=1, subsample=0.9. The consensus LCL dataset and the script used for CENTRE final training can be found at: http://owww.molgen.mpg.de/~CENTRE_data/CENTRE_final_training.zip

#### Comparison with TargetFinder

CENTRE features were calculated based on the ENCODE Registry of cCREs on hg38 and the basic gene annotation of GENCODE Release 40 (GRCh38). However for the comparison with TargetFinder on the BENGI datasets we used the hg19 annotation of cCREs-ELS and GENCODE v19 TSS that the authors originally used. CENTRE feature values were averaged for hg38 ET pairs corresponding to the same hg19 pair.

#### TargetFinder

We reimplemented TargetFinder (Whalen *et al*., 2016) (GradientBoostingClassifier, n_estimators = 4000, learning_rate = 0.1, max_depth = 5, max_features = ‘log2’, random_state = 0) to run on the BENGI ET pairs with the customized CV scheme suggested in (Moore *et al*., 2020). We calculated all the 303 features on LCLs, Hela, K562, IMR90, and NHEK datasets using the generate_training.py script and the corresponding datasets provided on the Github page (https://github.com/shwhalen/targetfinder).

Regarding the five tissue datasets where not all genomic datasets required by the TargetFinder method were available, we implemented a TargetFinder reduced model using a subset of 13 features coming from DNase, H3K4me3, H3K27ac, and CTCF experimental assays and distance. We used experimental assays downloaded from the ENCODE portal (Supplementary Table 5). Genomic features for all the implementations were calculated for enhancer, promoter, and window regions (EPW setting).

## Code Availability

CENTRE R software is accessible via https://github.com/slrvv/CENTRE

## Acknowledgements

Funding from German Ministry of Education and Research (BMBF, FKZ 01IS18037G] is gratefully acknowledged.

